# Phase transition in atomistic simulations of model membrane with thylakoid lipids of red algae

**DOI:** 10.1101/2022.10.19.512972

**Authors:** Arun K. Rathod, Dhruvil Chavda, Moutusi Manna

**Affiliations:** Applied Phycology and Biotechnology Division, CSIR Central Salt & Marine Chemicals Research Institute, Bhavnagar 364002, Gujarat, India

**Author notes:** Moutusi Manna **Email:**. **Author Contributions:** MM: Conceptualization and design of the study, Analysis and interpretation of data, Manuscript writing; AKR: Performed system preparation, simulations, data acquisition and analysis; DC: Performed data analysis. **Competing Interest Statement:** The authors declare no competing financial interest.

**Keywords:** Temperature, lipid phases, domain coexistence, lipid segregation, fatty acid unsaturation

## Abstract

Marine algae are diverse photosynthetic organisms, profoundly rich in bioactive compounds. Temperature is a major factor in algal cultivation and biomass production. At the cellular level, the change of temperature is reflected in oscillating algal lipid/fatty acid profile and inhibition of photosynthetic activities. The function of thylakoid membrane system is intimately dependent on its lipid matrix, however the molecular organization of these lipid membranes and particularly their adaptive arrangements under temperature stress remain largely unexplored. The present work employing extensive atomistic simulations provides the first atomistic view of the phase transition and domain coexistence in model membrane composed of thylakoid lipids of a marine alga, between 10-40 °C. The model membrane undergoes a transition from a gel-like phase at 10-15 °C to a homogeneous liquid-disordered phase at 40 °C. Clear evidences of spontaneous phase separation into coexisting nanoscale domains are detected at intermediate temperatures. Particularly at 25-30 °C, we identified the formation of a stable rippled phase, where the gel-like domains rich in saturated and nearly hexagonally packed lipids separated from fluid-like domains enriched in lipids containing polyunsaturated chains. Cholesterol impairs the phase transition and the emergence of domains, and induces a fairly uniform liquid-ordered phase in the membrane over the temperatures studied. The results have implications in understanding the role of lipids in temperature adaptation in algal.

## Introduction

Marine macroalgae are diverse photosynthetic organisms, increasingly viewed as potential renewable sources of bioactive compounds and nutritionally beneficial polyunsaturated fatty acids (PUFAs) (1, 2). Red algae are one of the oldest and largest groups of marine algae. They are long been commercially exploited as a source of hydrocolloids, like agar and carrageenan, widely used in food industries (3). Compared to terrestrial plants, marine algae are exposed to diverse environmental stress including fluctuations in temperature, salinity, nutrients and light (4). Temperature is a major factor that varies across the seasons, location, day-night cycle, etc. Alteration in lipid composition and degree of acyl chain unsaturation is a well-documented phenomenon in algal response to high and low temperatures (5-9). By adjusting membrane constituents cells maintain the fluidity or phase stability of their cell membranes for optimal functioning, in a process called “homeoviscous adaptation” (5, 10). Photosynthesis is one of the most sensitive processes determining algal response to temperature stress and lipids are believed to play a key role in the algal acclimation process (11, 12).

The light reactions of photosynthesis take place at thylakoid membranes. These are the extensively developed internal membrane systems of the chloroplast and the main intracellular membranes in algae (13, 14). The bulk of the membranes are constituted by lipids that house numerous proteins, protein complexes, and cofactors; all work in coordination to harness the energy of light. The lipid bilayer of thylakoid membranes is mainly composed of four unique lipid classes: monogalactosyldiacylglycerol (MGDG), digalactosyldiacylglycerol (DGDG), sulfoquinovosyldiacylglycerol (SQDG), and phosphatidylglycerol (PG) (Fig. S1), each has specific roles in the maintenance of thylakoid membranes and photosynthesis. The composition is conserved across organisms and distinctly different from other biological membranes (13-17).

Thylakoid lipids are extensively studied for the propensity to form non-bilayer phases (18- 20). MGDG favors the formation of inverted hexagonal or cubic phases when dispersed in water, whereas DGDG, SQDG, and PG prefer the conventional lipid bilayer structure. The coexistence of bilayer and non-bilayer phases in isolated thylakoid membranes is identified by several studies (18, 20). In contrast, other studies show that the non-bilayer structures are not observed unless the membrane is stressed (21, 22). *In vivo* thylakoid membranes are organized as bilayers. Membrane-spanning proteins are hypothesized to be crucial in imposing the bilayer configuration (18). A higher MGDG/DGDG ratio promotes non-bilayer phases (18, 19). However MGDG mixed with other thylakoid lipids at ratio observed *in vivo* can produce bilayer structure without proteins (22). There is a lack of consensus regarding the lateral heterogeneity in thylakoid membranes. The formation of domains has been reported and the different regions of these membranes are thought to have different lipid compositions (23-27). Whereas other studies suggest that the bulk lipids of thylakoid membranes do not display lateral heterogeneity (28). The experimental techniques are limited to mesoscopic scale resolution and the nanoscale organization of thylakoid membranes remains largely unexplored.

Ample evidence indicates that the fluidity of thylakoid membranes may play an important role in controlling photosynthesis (29, 30). For example, the unsaturation of thylakoid lipids potentially stabilizes the photosynthetic machinery against low-temperature photoinhibition (31, 32). While the exposure of isolated chloroplast to short-term heat stress is known to alter the structural organization of thylakoid membranes and photosynthetic activity (30, 33). Although sterols are not present in thylakoids in detectable amounts, artificial modulation of membrane fluidity by cholesterol and cholesterol-analogues are reported to affect the rate of electron transport, proton uptake, and lateral diffusivity of cofactor and protein-pigment complexes (29, 30, 34, 35). While the link between the fluidity and functionality of thylakoid membranes is established, the mechanism is far from being completely understood.

Membrane fluidity is greatly influenced by temperature. For a one-component lipid bilayer, as the temperature drops below the phase transition temperature, the membrane changes from a disordered liquid state (liquid-disordered phase, L_α_) to a highly ordered solid state (gel phase, L_β_) (36). An intermediate and comparatively less-studied “rippled phase” (P_β_) phase may exist for certain lipids, in which L_β_-/L_α_-like regions coexist (37, 38). Cholesterol regulates the fluidity of the membrane. When added in low concentrations, it increases the acyl chain order of fluid-phase saturated lipids, whereas at high concentrations it induces a liquid-ordered (L_o_) phase (39, 40). In model membranes, coexisting liquid phases (L_o_/L_α_) have been observed in a wide range of compositions and temperatures (41-43). Cholesterol plays a key role in domain formation. According to the “raft model”, plasma membrane includes nanoscale functional domains enriched in cholesterol and sphingolipids. Nanodomains are known to be vital for many biological processes including signal transduction, trafficking, and membrane protein function (42, 43). However, due to small size and dynamic nature, the detailed molecular structures of the ordered domains are scare experimentally.

Molecular dynamics (MD) simulations have been extensively employed to study lipid membranes and membrane proteins (44-49). Phase properties of simplified model membranes are accurately predict via computer simulations (39, 43). Several MD simulations have investigated phase separation in membranes at the coarse-grained level, (10, 50) but more accurate and computationally demanding all-atom simulations are limited (42). To bypass the long time scale required to capture spontaneous phase separation, atomistic simulations are often performed with preformed coexisting domains (51-52). Computational studies enhance our understanding of the physical principles underlying nanodomains formation and provide a detailed molecular view of nanodomains that are otherwise inaccessible from experiments (42, 43, 51, 52). Bilayers containing an unsaturated lipid (*e*.*g*. dioleoylphosphatidylcholine, DOPC), a saturated lipid (*e*.*g*. dipalmitoylphosphatidylcholine, DPPC, or sphingomyelin), and cholesterol are frequently simulated to study domain-coexistence, as a canonical model of the plasma membrane (41-43, 50-52). However, the lipid composition of thylakoid membranes is distinctly different. They are mainly composed of noncommon galactolipids and sulfolipids but are largely devoid of cholesterol (13-17). The greater acyl chain variety of thylakoid lipids and richness in PUFAs further adds to the compositional complexity (14). So far only a few computational studies have focused on thylakoid membranes (53-55). A previous computational study has compared the properties of thylakoid membranes from cyanobacteria and a higher plant at their respective physiological temperatures (53). Others include very short all-atom simulations of photosystem II (PSII) in the thylakoid membrane (54, 55).

The present work employs extensive atomistic simulation (totaling ∼55 µs) to study the phase behavior of a model thylakoid membrane based on the experimentally determined lipid and fatty acid composition of a commercially important marine alga *Gracilaria Corticata*, collected from the Veraval coast of the Saurashtra region in India (56, 57). The seawater temperature in this region varies between 20-32 °C across the year and ∼25-28 °C during the algal growth period (58). The alga exhibits a high DGDG to MGDG ratio and has saturated palmitic acid (C16:0) and polyunsaturated arachidonic acid (C20:4 ω-6) as predominant fatty acids and oleic acid (C18:10 as main monounsaturated fatty acid (MUFA) (56, 57). We model a complex bilayer consisting of polar heads MGDG/DGDG/SQDG/PG in the ratio 1.28/2.25/1/1.46, in fair agreement with experiments (57). The acyl chain combinations used here are: di-palmitoyl, palmitoyl-oleoyl, palmitoyl-arachidonoyl, with a ratio of total saturated fatty acid (SFA)/MUFA/PUFA equals 3.52/1/2.59 that closely mimics experimental value (56). Simulations are conducted over the temperature range 10-40 °C to study the properties of the model algal membrane at physiological temperatures and under cold and heat stress. We investigated the structure and physical properties of resulting lipid phases and domains and the driving forces for domain formation in the model thylakoid membrane (referred to as Thylakoid-LBM), starting from a random conformation. We also studied the effect of cholesterol (25 mol%) on phase states and the emergence of domains in the cholesterol-treated thylakoid membrane model, referred to as Chol-Thylakoid- LBM. Although cholesterol is not found in thylakoids, it is the dominant sterol in red algae (5) and is often treated as a well-known modulator of membrane fluidity. Our simulations show for the first time the phase transition and phase separation in a model thylakoid membrane as a function of temperature and the phase stability induced by cholesterol. To the best of our knowledge, these are the first simulations model focused on cell membranes of marine macroalgae.

## Results

### Phase transition in the model membrane with thylakoid lipids

The most prominent observation in the present study is the phase transition of the model Thylakoid-LBM membrane as a function of temperature. As a criterion for defining membrane phases (gel, L_o_, and L_α_ phases) here we determine the lipid hydrocarbon chain order parameter (-S_CD_) as widely used in earlier studies (39, 42, 43). Fig. 1 presents the -S_CD_ distribution along the *sn-1* palmitoyl chain over all lipids and the average -S_CD_ over all carbons in that chain, while the -S_CD_ of other acyl chain types are included in the supporting information (Fig. S2). Although the experimental data on the specific thylakoid lipid mixture or lipid types are not available for comparison, our results are compared against and are in excellent agreement with the -S_CD_ data of the palmitoyl chain of DPPC lipid in various phases derived from previous Nuclear Magnetic Resonance (NMR) measurements and simulations (36,40,43, 51).

**Fig. 1.**
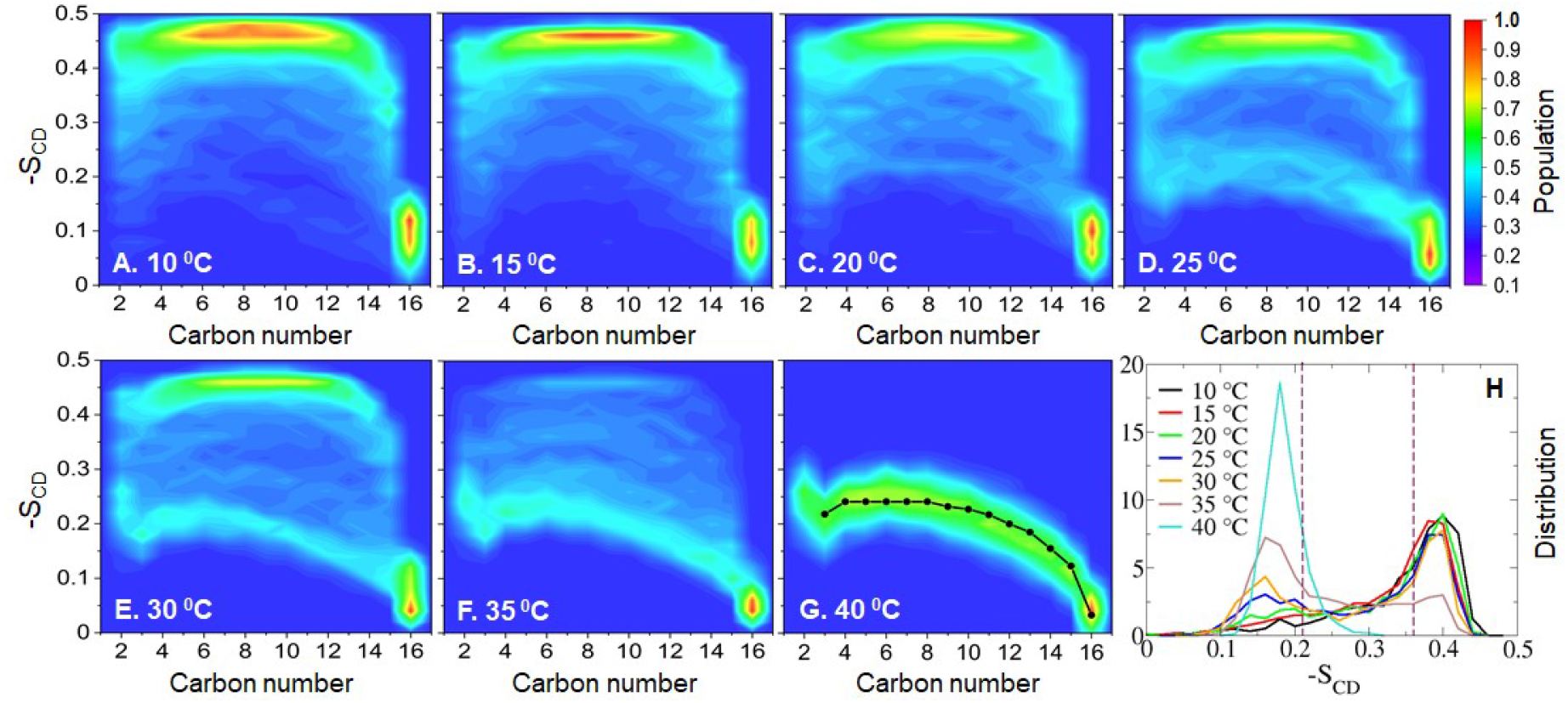
Lipid acyl tail order parameter (-S_CD_) of the thylakoid membrane model as a function of temperature. (A-G) -S_CD_ distribution along carbon position of the *sn-1* palmitoyl chain. The available experimental -S_CD_ profile (36) is shown as black line with dots. (H) Distribution of average -S_CD_ over all carbon atoms of the *sn- 1* chain. Available experimental data on L_o_ and L_α_ phases from ref (51) are shown as dotted lines at 0.36 and 0.21 respectively.

As shown in Fig. 1, at the lowest temperatures, 10-15 ºC, the lipid acyl chains are highly ordered. No appreciable increase in order is noted upon cooling from 15 ºC to 10 ºC and the phase properties at these two temperatures are comparable. Average -S_CD_ distributions show a peak at 0.4 (Fig. 1H). The value is higher than the NMR-derived value of 0.36 on the L_o_ phase (51). Further, the -S_CD_ distribution along the palmitoyl chain obtained from our simulations is in good agreement with that in the gel phase reported in all-atom simulations (43). These results suggest that the model membrane studied here is in a gel-like state at 10-15 ºC. However, we noticed that even at this low-temperature range the membrane is not completely homogeneously ordered, and contains minor proportions of disordered regions (average -S_CD_ < 0.3, Fig. 1H). On the other hand, at the highest temperature, 40 ºC, the model membrane is most disordered and shows a narrow average -S_CD_ distribution, in line with the behavior of a homogeneous phase, with a peak at 0.18 (Fig. 1H). The -S_CD_ distribution along the *sn-1* chain is also indicative of the homogeneous L_α_ phase at 40 ºC (Fig. 1G). Our result is fairly close to the average -S_CD_ (0.21) reported by NMR (51) and nicely overlaps with the experimental -S_CD_ profile of DPPC in the L_α_ phase at 41 ºC (36).

Notably, between 20 ºC to 35 ºC, the model membrane undergoes a phase separation. The two distinct peaks of average -S_CD_ at ∼0.4 and ∼0.16, imply the coexistence of ordered and disordered domains, respectively (Fig. 1H). The -S_CD_ distributions along the *sn-1* chain are significantly broad and show contributions from both the highly ordered gel-like and disordered fluid-like L_α_ phases (Fig. 1), as we observe at low and high temperatures respectively. Particularly, the distributions at 25 ºC and 30 ºC show two distinct clusters appear like a superposition of two separate -S_CD_ profiles from gel-like and L_α_ phases (Fig. 1D-E). Contribution from liquid-ordered (L_o_) phase seems to be absent or less significant in the -S_CD_ distributions, but observed in presence of cholesterol as described in detail below. A larger fraction of gel-like character is detected at the lower temperature, while the fluid-like nature is more pronounced at 35 ºC. Clearly, the proportion of disordered region gradually increases with temperature; until the membrane transforms to a homogeneous L_α_ phase at 40 ºC (Fig. 1G). These results, therefore, suggest a gel-to-L_α_ phase transition in the model membrane at 10-40 ºC.

The phase transition is also evident from the change in other structural properties, like area-per-lipid and thickness of the model membrane. As expected the area-per-lipid grows and consequently the thickness decreases with temperature (Fig. S3 and S4). Between 20-30 ºC these properties show an intermediate behavior (Fig. S4). The area-per-lipid substantially decreases and the membrane becomes more densely packed at 15 ºC. No significant condensation in the area is observed upon further cooling down to 10 ºC, which is consistent with the gel-like behavior at low temperature. While a sharp rise in area-per-lipid occurred at 35 ºC. At 40 ºC the membrane is most diffused supporting its disordered fluid-like character while still preserving the lamellar structure.

Changes in the lipid phase as a function of temperature also affect the interactions of the membrane with ions and waters (Fig. S5). Due to the presence of anionic lipids PGs and SQDGs, the membrane is negatively charged and it exhibits a clear selectivity towards K^+^ ions over Cl^−^ ions. Interestingly with an increase in temperature, the number of lipid-K^+^ contacts decreases with a parallel increase in lipid-Cl^−^ contacts. The number of lipid-water contacts gradually increases with temperature and a sharp rise is noted at 35 ºC and above, when the membrane is highly diffused.

### Molecular structures of lipid phase and nanodomains in the model thylakoid membrane

Let us now look into the detailed molecular structure of the various lipid phase and nanoscale domains formed in the model thylakoid membrane (Fig. 2, 3, and S6). Selected snapshots in Fig. 2 show a very dense packing of lipids at 10 ºC. The lipid acyl chains are most ordered and neatly arranged, as typically found in the gel-like domains. The tilt angle distribution of the *sn-1* chains shows that the chains are oriented nearly parallel (<10°) to the bilayer normal (Fig. S7A), in excellent agreement with recently reported simulations on gel-phase DPPC/DOPC bilayer (42). Earlier simulations on gel phase in one-component membranes have reported tilting of highly condensed saturated lipid chains, in order to adapt to the larger lipid headgroups (38, 39). Such tilting of tails is apparently not required in our case as the presence of lipids with unsaturation in the model membrane already resulted in a larger average area per lipid chain. The top view of the snapshot at 10 ºC also reveals the gel-like packing of the lipid hydrocarbon chains (Fig. 3A). Another interesting feature clearly evident from the snapshots (Fig. 3A and 3D) is the nice arrangement of lipid hydrocarbon chains in a hexagonal pattern almost throughout the membrane plane, except for small patches with distorted packing. The nearest neighbor distributions of lipid chains, Fig. 3H, show a peak at six, which indicates that a majority of the lipids at 10-15 ºC have six nearest neighbors, supporting the hexagonal packing. Similar hexagonal packing was detected within the L_o_ phase by NMR and simulations (43, 51, 52). Further, to examine the spatial distribution, we calculated the thickness map (Fig. 3I-K and S6I-L) and “jump-map” (Fig. S8), where the latter describes the average in-plane lipid displacements calculated over a 1 ns interval. Consistent with the lipid chain orientation and order, the membrane has the highest hydrophobic thickness and the lowest lipid mobility at low temperatures (Fig. S8).

**Fig. 2.**
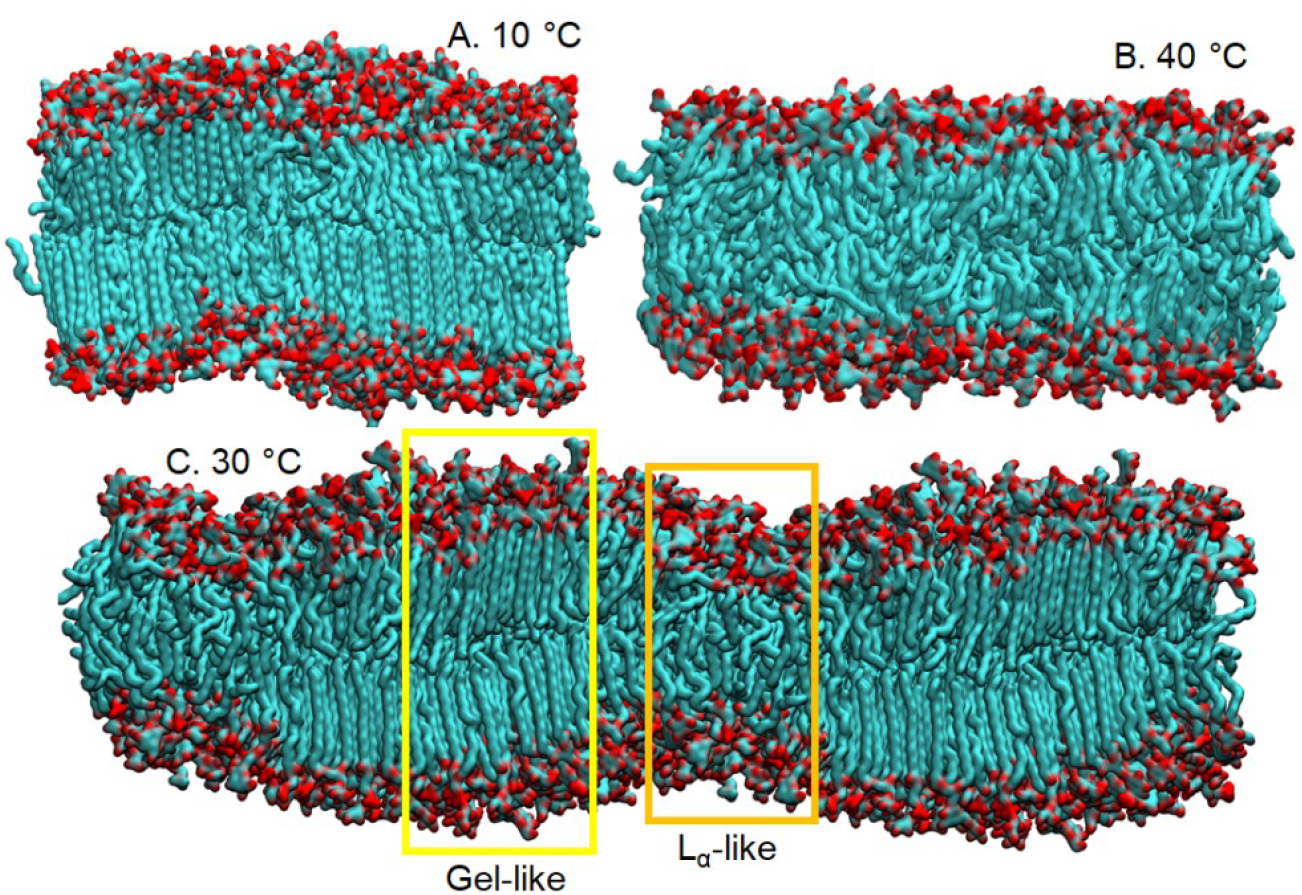
Representative snapshots of the model thylakoid membrane in: (A) a gel-like phase at 10 °C and (B) a L_α_ phase at 40 °C. (C) Snapshot with its periodic image shows the gel-L_α_ coexistence in the membrane with a wave-like surface at 30 °C. Lipid carbon atoms are colored in cyan and other non-hydrogen atoms in red. Waters and ions are not shown for clarity.

**Fig. 3.**
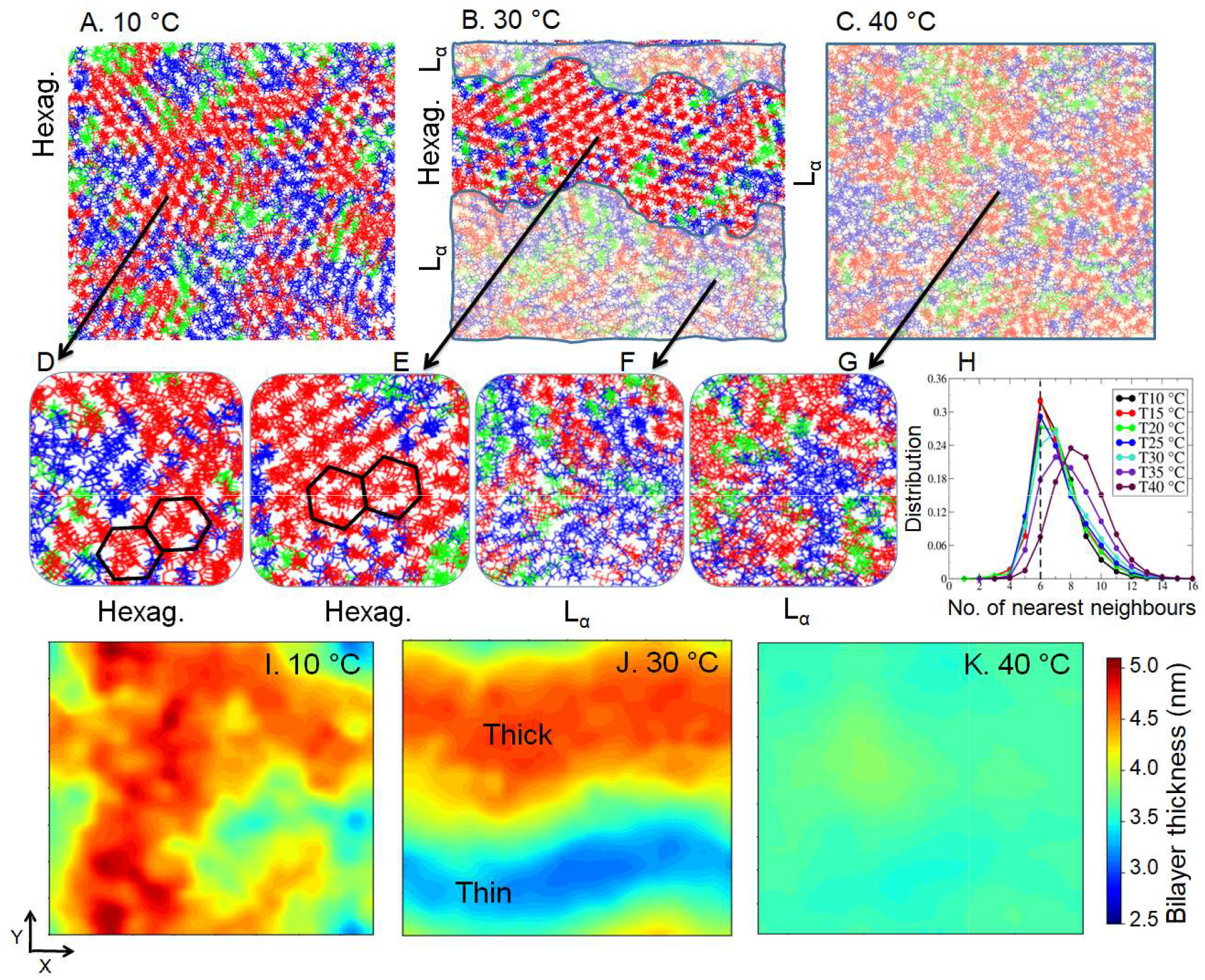
(A-C) Top-views of the thylakoid-LBM, and (D-G) the zoomed images at different temperatures. Lipids with di-palmitoyl, palmitoyl-oleoyl and palmitoyl-arachidonoyl chains are colored in red, green, and blue, respectively. Shaded areas in the snapshots (B, C) represent L_α_-like regions with loosely packed lipids (F, G). Unshaded areas the snapshots (A, B) represent gel-like regions with app oximately hexagonally packed lipids (D, E). (H) The number of nearest neighbour distribution of lipid hydrocarbon chains. (I-K) Bilayer thickness (nm) projected on membrane plane.

On the contrary, the snapshot (Fig. 3C) at 40 ºC shows a nearly uniform lipid distribution. It is readily apparent from the snapshots that the lipid chains are highly disordered and splayed (Fig. 2B). The distribution of acyl chain angle is significantly broad with a peak near ∼20º (Fig. S7A). Similar behavior was reported by previous simulations on fluid phase lipid membranes (42). The hexagonal packing of lipid acyl chains is disrupted (Fig. 3G) and lipids are shown to have more, ∼8-9 nearest neighbours (Fig. 3H). The nearest neighbour distribution became much broader at the elevated temperature with a clear shift in peak position (Fig. 3H). At 40 ºC the thickness of the bilayer is ∼3.5-3.7 nm, which is the lowest of all temperatures and strikingly less than that we obtained for the gel-like domains (Fig. 3). Further, the thickness map at 40 ºC is fairly uniform, suggesting a homogeneous L_α_ phase. The mobility of the lipids in the L_α_ phase is ∼5 times faster than observed for the gel-like phase (Fig. S8). These properties may again suggest the gel-to-fluid transition of the model thylakoid membrane at the temperature range simulated here.

The co-existence of ordered and disordered domains in the model thylakoid membrane is noted between 20 ºC to 35 ºC temperatures (Fig. 2, 3, and S6). We observed spontaneous phase separation of lipids into domains with distinctly different properties. Depending on the temperature the domains are either stable or dynamic. The heterogeneity is validated by the thickness map and jump maps (Fig. 3I-K, S6I-L, and S8). For example, the thickness plot at 30 ºC shows the clear formation of alternative thick and thin domains in the model membrane (Fig. 3J), a typical signature of rippled phases (37, 38). Such a large difference (∼1.5 nm) in thickness gives rise to the wave-like surface (Fig. 2C). The thick domains, commonly termed as “major arms” corresponds to the gel-like component with an average -S_CD_ of ∼0.4 (Fig. 2C, 3B,E, and 1H). The tightly packed nature of these domains leads to very slow, collective dynamics, as reflected from the lipid diffusivity, which is much slower than the thinner domains (Fig. S8E). The thin domains, which are known as “minor arms”, on the other hand, correspond to the fluid-like component with highly disordered acyl chains (average -S_CD_ ∼0.16) (Fig. 2C, 3B,F, and 1H). The kink regions are formed with ordered lipid chains in one but disordered chains in the opposing leaflet (Fig. 2C). In line with phase separation at 30 °C, the top view of the system shows spontaneous segregation of lipids into different domains (Fig. 3B,E,F). It is evident from the snapshot that the fully saturated lipids are partitioned more favorably into the ordered domains (Fig. 3B,E). Enrichment of saturated chains is associated with the formation of a longer-range hexagonal order (Fig. 3B,E,H). Hexagonal substructures are formed mainly by the lipids with both saturated tails. Lipids with unsaturated chains, particularly PUFAs, are largely excluded from the ordered domains and segregated more into the disordered domains (Fig. 3B,E,F). Because of the higher concentrations of unsaturated lipids, these disordered domains are even less ordered (average -S_CD_ ∼0.16) than the homogeneous L_α_ phase (average -S_CD_ ∼0.18) at high temperatures. Demixing of lipids is clearly depicted in Fig. S9A. At 30 °C, We found ∼68% of fully saturated lipids in the ordered but only ∼32% in the disordered regions, while out of PUFA-containing lipids ∼68% are detected in the disordered region and only ∼32% in the ordered region. Interestingly, the polar lipid head groups exhibit no clear domain preferences and are almost uniformly distributed between coexisting domains (Fig. S9B). A stable gel-like/fluid-like domain coexistence and formation of local hexagonal structure in the ordered domain are also observed at 25 ºC (Fig. S6C,G,K). Phase separation occurs, but to a lesser extent at 20 ºC. While at 35 ºC, lipids are highly dynamic (Fig. S8F) and the heterogeneous features of the membrane are not sufficiently captured by the time-averaged analyses employed here.

### Cholesterol induces and stabilizes the liquid-ordered phase over the temperature range

Let us now turn to the effect of 25 mol% cholesterol on the phase behavior model thylakoid membrane. Our results reproduced the well-known condensation effect of cholesterol on lipid membranes (39, 59). Fig. S4 convincingly illustrates the condensing effect, the area per lipid molecule decreases at all temperatures in presence of cholesterol, more than that would be expected on the basis of ideal mixing. Consequently thickness of the bilayer increases. Cholesterol also reduces the number of lipid-water and lipid-ion contacts (Fig. S5), supporting the view that it may protect the membrane from being too leaky at elevated temperatures.

Cholesterol strongly modifies the membrane phase at all temperatures (Fig. 4, S10 and S11). A notable impact is observed between 20-30 °C. Including cholesterol into the thylakoid lipid mixture, transforms the rippled phase as observed in absence of cholesterol (Fig. 1C-F), into a liquid-ordered (L_o_) phase (Fig. 4B). The -S_CD_ distribution is much narrower in presence of cholesterol, suggesting the homogeneous phase behavior (Fig. 4 and S11). The lipids acyl chains are ordered, as also seen from the snapshots, and have ordered parameters slightly less than gel-phase but much higher than the L_α_ phase. Such properties are typical for L_o_ phase. -S_CD_ profile at 25 °C (Fig. S11C) fairly overlaps with the profile derived from NMR for DPPC/cholesterol mixture in pure L_o_ phase at 25 °C (40). The average -S_CD_ distribution shows a peak at 0.36, in total agreement with the experimental data (0.36) on L_o_ phase of DPPC/DOPC/cholesterol mixture, (51) further supporting the liquid-ordered nature. Our results suggest that cholesterol promotes the ordered-like structure, while the gel-like contribution decreases, consistent with previous findings (39, 42).

**Fig. 4.**
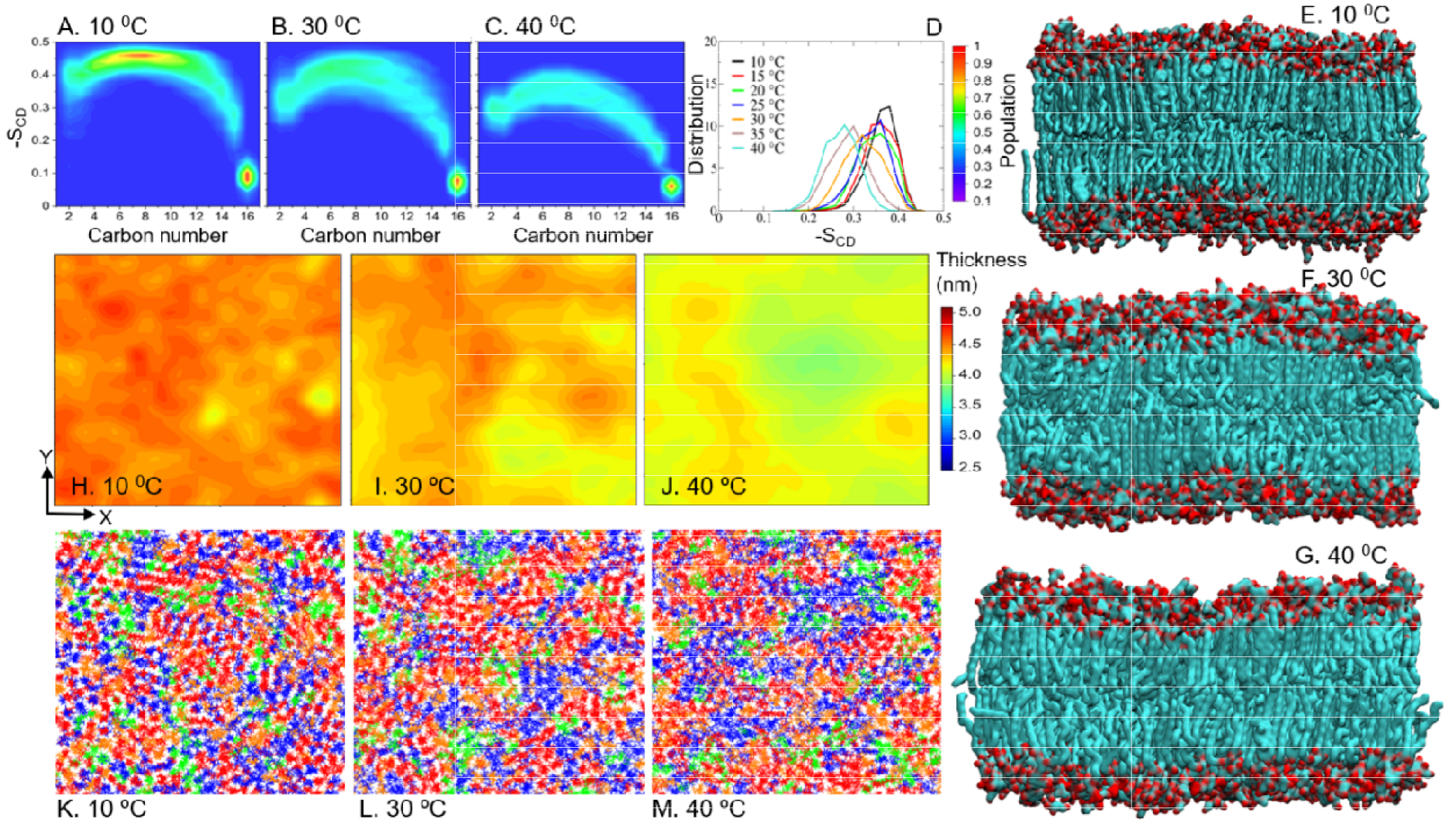
Properties of Chol-Thylakoid-LBM. (A-G) -S_CD_ distribution along carbon position of the *sn- 1* palmitoyl chain. (D) Distribution of average -S_CD_ over all carbon atoms of the *sn-1* chain. (E-G) Representative snapshots of the side-view of the system which same color code as Fig. 2. (H-J) Thickness maps at different temperatures. (K-M) Top-view of the membrane showing the lateral organization of lipids at different temperatures. Lipids with di-palmitoyl, palmitoyl-oleoyl and palmitoyl-arachidonoyl chains are colored in red, green, and blue, respectively. Cholesterols are shown in orange.

We found that cholesterol also significantly modifies the membrane properties at high temperatures. Including cholesterol transforms the L_α_ phase developed in absence of cholesterol (Fig. 1G), into L phase (Fig. 4C). With the rise in temperature, the order parameter slightly decreases in cholesterol-containing membranes (Fig. 4 and S11A-D), which is however nowhere close to the values obtained in absence of cholesterol. Even at the highest temperature, 40 °C, the membrane with cholesterol is significantly more ordered. Our results show that at high temperatures cholesterol makes the membrane less fluid. While at low-temperature addition of cholesterol increases the fluidity and transforms the gel phase into L_o_ phase. Only at the lowest temperature simulated here, *i*.*e*. at 10 °C, the -S_CD_ values are close to the gel phase (average - S_CD_ 0.4) (Fig. 4A). Among the different acyl tail types used in the present study, cholesterol regulates the ordering of the saturated chains the most, followed by the mono-unsaturated chains (Fig. S10). The PUFA chains are least affected by cholesterol, which is again in fair agreement with earlier studies (60). Overall, our results show that including cholesterol into the thylakoid lipid mixture significantly stabilizes the phase state and inhibits gel-to-fluid transformation over the temperature range studied.

In presence of cholesterol nearly homogeneous lipid distributions are identified at all temperatures (Fig. 4K-M and S11I-L). As apparent from the snapshots cholesterol limits the large-scale phase separation and increase the miscibility of lipids. Fig. S12 shows that cholesterol almost equally likely interacts with both fully saturated lipids and PUFA-containing lipids up to 30 °C. The results are in line with the previous studies showing that cholesterol, by virtue of two different faces of its steroid ring - a smooth face and a rough face with methyl groups, enhances the lateral mixing amongst the saturated and unsaturated lipids (42, 43, 51). In addition, we found that cholesterol also perturbs the long-scale hexagonal packing of lipid hydrocarbon chains (Fig. 4). The spatial homogeneity of the membrane in presence of cholesterol is further reflected from the thickness-map (Fig. 4 and S11) and the jump-map (Fig. S13); these properties show intermediate values between the gel and L_α_ phases observed in absence of cholesterol.

## Discussion

Increasing evidence convincingly shows the effect of temperature on the functioning of the thylakoid membrane systems (29-35). These effects are thought to be attributed mainly to the temperature-induced modulation of thylakoid membrane fluidity, which may affect the lateral separation of the main pigment-protein complex, mobility of electron carrier for the electron transport reactions, physical movement of light-harvesting chlorophyll *a/b* complex for energy distribution between PSI and PSII, among others. Several studies have reported that algal lipid and fatty acid composition is highly sensitive to ambient temperature (5-9). Changes in the lipid unsaturation level inevitably reflect membrane fluidity. It is believed that cell membranes utilize lipids to maintain the structural integrity, selective permeability as well as stability of the photosynthesis apparatus. To get a deeper understanding of the mechanism of stress tolerance, it is important to understand how the thylakoid lipid membrane changes its physical properties and phase states to adapt to the fluctuating temperature. The previous studies on algal lipids mostly encompass the elucidation of lipids and their fatty acid profiles. To the best of our knowledge, no study so far has characterized the physical properties of algal thylakoid membranes under adaptive rearrangements.

The present work employs robust atomistic simulations (totaling ∼55 µs) to characterize the effect of temperature on the phase behavior of the thylakoid membrane model of the red alga *G. Corticata*. We present here the first atomistic view of phase transition and phase coexistence in the algal thylakoid membrane model over the temperature range 10-40 °C. It is observed that at the highest temperature membrane is in a homogeneous liquid-disordered (L_α_) phase. At low temperatures, the membrane has typical characteristics of a gel phase, in which the lipid hydrocarbon chains are packed tightly and organized laterally in an approximately hexagonal structure. The global mixing of lipids is rather homogenous although small patches of disordered regions do seem to occur at low temperatures. At intermediate temperatures, the membrane undergoes spontaneous phase separation. At 25 °C and 30 °C, formation of stable rippled phase is detected, where the gel-like domains, in which the bilayer is thick and lipids ordered, are separated from the fluid-like domains, in which the bilayer is thin and lipids are disordered. Our simulations start with random lipid mixture and co-existing domains developed over the course of microseconds long atomistic simulations and independent trajectories. Noticeably, the structural and dynamical properties studied here point that the membrane nearly sustains its properties between 20-30 °C that vary appreciably at low and high temperatures. Despite of the complexity of the thylakoid lipid mixture, many of the current observations match the known phase diagram data of pure lipid or binary lipid mixtures like DPPC/DOPC, (38, 39, 42, 43) which serve to validate our results.

The scope for elucidation of phase properties and the structure of nanodomains in native thylakoid lipid membranes is very limited. However, of direct relevance to our finding, a previous study using freeze-fracture electron microscopy had captured the formation of gel phase lipid at 15 °C in the thylakoid membrane of thermophilic cyanobacteria that grow at 38 °C (23, 25). In addition, this study also showed the presence of smooth particle-free patches in these membranes that signals the presence of phase-separated gel phase lipids from which intrinsic proteins are excluded. Further, the formation of gel-phase lipids was confirmed by X-ray diffraction measurements (26). These studies not only support the phase separation into coexisting domains in thylakoid membranes in line with that found in our simulations but also suggest the potential role of such domains in controlling the partitioning of integral membrane proteins (23-26). It is worth mentioning the red alga *G. Corticata* studied here, abundantly grows in the Saurashtra coast of India during autumn-winter-spring when the seawater temperature ranges ∼25-28 °C (58). Interestingly, the stable ordered/disordered coexisting domains in the thylakoid membrane model are also most prominently seen in our simulations at 25 °C and 30 °C, further suggesting that such membrane domains might be of functional relevance. Functional nanoscale domains in the plasma membranes have been extensively studied owing to their role in many vital cellular processes. However, the role of such domains in the thylakoid membrane and their relation to photosynthesis is yet to be established.

Our results highlight the importance of fatty acids as a key player in the modulation of nanodomain formation in the thylakoid membrane model. Several earlier experimental studies have focused on the role of polar lipid headgroups, particularly MGDG in bilayer to non-bilayer phase transition and in the structure and dynamics of the chloroplast thylakoid membrane (14, 16, 18, 19). The present simulations show that the model thylakoid membrane of this specific red alga, which has more DGDG than MDGD, sustains its lamellar structure through the simulated temperature range of 10-40 °C. Lipid polar headgroups show no such apparent domain preference, and their global mixing is rather homogeneous, which is in agreement with previous studies (28, 53). Whereas we found that the nanodomain formation is driven by preferential segregation of lipids into differentially ordered domains depending on the lipid acyl chain types. The saturated lipids are preferentially segregated into ordered regions. Among the various lipid acyl chains present in the system, polyunsaturated chains are least affected by temperature and enrichment of PUFA chains in the disordered domains further suggests that an increase in their content may protect the membrane from low-temperature rigidification. Indeed, marine macroalgae from cold-water environments are found to have higher PUFA contents than macroalgae from warmer waters, (6-8) while the thermophilic cyanobacteria that grow at high temperature, contain more saturated lipids (17). Our results provide a plausible explanation to a vast experimental observation showing the alteration in fatty acid unsaturation as the most common change in algae under temperature stress.

We have also studied the effects of cholesterol on the thylakoid lipid matrix as a function of temperature. Several earlier studies have shown that introducing cholesterol or its analogue into the thylakoid lipid phase is accompanied with inhibition of electron transport, changes in ionic conductivity and fluorescence (29, 30, 34, 35). These cholesterol-induced effects on the functioning of photosynthetic processes are proposed to be associated with the ability of cholesterol or sterols in general, to modulate the fluidity of lipid membranes. Our simulations provide the first insights into how cholesterol regulates the phase states of the thylakoid lipid membrane with atomistic details. Our results suggest that cholesterol stabilizes the model thylakoid membrane both from rigidification at low temperature and being too fluid and leaky at high temperature. Cholesterol induces and stabilizes a fairly homogeneous L_o_ phase in the membrane over the temperatures studied here. Cholesterol also significantly suppresses the phase-separation and emergence of domains in the model membrane and increases lipid miscibility. Many of these cholesterol-induced effects are consistent with experiments and simulations on natural and artificial membranes. Here we used cholesterol as an agent to modify lipid microviscosity; as it is widely used in animal and artificial membranes for the same purpose (39, 40, 43, 59, 60). However, it might be worth exploring in the future more physiological options, like carotenoids or α-tocopherol, to carefully regulate the properties of the thylakoid lipid membrane (61, 62) The phase stability of thylakoid lipid membranes is important to stabilize the photosynthetic apparatus under temperature stress and is particularly relevant in the context of global warming. An increase in seawater temperature affects the cultivation of marine algae. It has been witnessed that even a sudden, few degree raise in seawater temperature from normal temperature was associated with the mass mortality of a marine alga *Kappaphycus alvarezii* and had adversely impacted the socio-economic scenario of the coastal region (63, 64).

Summarizing, the present work provides the first atomistic insights of the gel-to-fluid phase transition and domain coexistence in a model algal thylakoid membrane between 10-40 °C. Our simulations reveal the previously unseen molecular structures of lipids domains spontaneously formed in the membrane starting from a completely random lipid organization. The incorporation of cholesterol induces a fairly uniform liquid-ordered phase and suppresses the phase transition and phase separation in the temperature range studied. The results improve our understanding of the lateral organization in thylakoid membrane with higher compositional complexity and have implications towards understanding the mechanism of lipid-dependent adaption of algae to temperature stress.

## Materials and Methods

We performed all-atom MD simulations of a thylakoid membrane model and the model treated with cholesterol at 7 temperatures from 10 °C to 40 °C, using CHARMM36 force field (65) and GROMACS 5.1.1 software package (66). Further details of lipid composition and methodology are given in SI.

## Supporting information

SI file includes methods and additional analysis.

## Acknowledgments

This work is performed using the high performance computer resources of CSIR Fourth Paradigm Institute, Bangalore, India. AKR and DC thank DBT for research fellowship. This manuscript has PRIS registration number 46/2021.

## Notes

### Competing Interest Statement

The authors have declared no competing interest.

## References

1. B. D. Elena-Suzana et al., Macroalgae—A sustainable source of chemical compounds with biological activities. Nutrients 12, 3085 (2020)

2. P. Kumari, M. Kumar, V. Gupta, CRK. Reddy, B. Jha, Tropical marine macroalgae as potential sources of nutritionally important PUFAs. Food Chem. 120, 749–757 (2010).

3. M. D. Torres, N. Flórez-Fernández, H. Domínguez, Integral utilization of red seaweed for bioactive production. Mar. Drugs 17, 314 (2019).

4. K. Mikami, S. Takio, Y. Y. Hiwatashi, M. Kumar, Environmental Stress-Promoting Responses in Algae. Front. Mar. Sci. DOI: 10.3389/fmars.2021.797613 (2021).

5. P. Kumari, M. Kumar, V. Gupta, CRK. Reddy, B. Jha, Algal lipids, fatty acids and sterols. Functional Ingredients from Algae for Foods and Nutraceuticals, 1st ed.; Herminia D., Ed.; Science Direct, 87–134 (2013).

6. E. Susanto, A. S. Fahmi, M. Hosokawa, K. Miyashita, Variation in lipid components from 15 species of tropical and temperate seaweeds. Mar. Drugs. 17, 630 (2019).

7. B. Narayan, K. Miyashita, M. Hosakawa, Comparative evaluation of fatty acid composition of different Sargassum (Fucales, Phaeophyta) species harvested from temperate and tropical waters. J. Aquat. Food Prod. Tech. 13, 53–70 (2005).

8. M. Nomura et al., Seasonal variations of total lipids, fatty acid composition, and fucoxanthin contents of Sargassum horneri (Turner) and Cystoseira hakodatensis (Yendo) from the northern seashore of Japan. J. Appl. Phycol. 25, 1159–1169 (2013).

9. M. Honya, T. Kinoshita, M. Ishikawa, H. Mori, K. Nisizawa, Seasonal variation in the lipid content of cultured Laminaria japonica: fatty acids, sterols, β-carotene and tocopherol. J. Appl. Phycol. 6, 25–29 (1994).

10. S. Erimban, S. Daschakraborty, Cryostabilization of the Cell Membrane of a Psychrotolerant Bacteria via Homeoviscous Adaptation. J. Phys. Chem. Lett. 11, 7709–7716 (2020).

11. R. Iglesias-Prieto, J. L. Matta, W. A. Robins, R. K. Trench, Photosynthetic response to elevated temperature in the symbiotic dinoflagellate Symbiodinium microadriaticum in culture. Proc. Natl. Acad. Sci. U.S.A. 89, 10302–10305 (1992).

12. D. Zou, K. Gao, The photosynthetic and respiratory responses to temperature and nitrogen supply in the marine green macroalga Ulva conglobata (Chlorophyta). Phycologia. 53, 86–94 (2014).

13. J. L. Harwood, Membrane lipids in algae. In Lipids in photosynthesis: structure, function and genetics. 1st ed.; Paul-André, S.; Norio, M., Ed.; Springer, Dordrecht 53–64 (1998).

14. J. Joyard, et al., Structure, distribution and biosynthesis of glycerolipids from higher plant chloroplasts. In Lipids in photosynthesis: structure, function and genetics. 1st ed.; Paul-André, S.; Norio, M., Ed.; Springer, Dordrecht 21–52 (1998).

15. Y. Nakamura, Plant phospholipid diversity: emerging functions in metabolism and protein– lipid interactions. Trends Plant Sci. 22, 1027–1040 (2017).

16. K. Kobayashi, H. Wada, Role of lipids in chloroplast biogenesis. In Lipids in Plant and Algae Development, 1st ed.; Nakamura, Y.; Li-Beisson, Y., Ed.; Springer, Cham 103–125 (2016).

17. I. Sakurai, et al., Lipids in oxygen-evolving photosystem II complexes of cyanobacteria and higher plants. J. Biochem. 140, 201–209 (2006).

18. W. P. Williams, P. J. Quinn, The phase behavior of lipids in photosynthetic membranes. J. Bioenerg. Biomembr. 19, 605–624 (1987).

19. G. Garab, B. Ughy, R. Goss, Role of MGDG and non-bilayer lipid phases in the structure and dynamics of chloroplast thylakoid membranes. In Lipids in Plant and Algae Development, 1st ed.; Nakamura, Y.; Li-Beisson, Y., Ed.; Springer, Cham 127–157 (2016).

20. C. Wilhelm, R. Goss, G. Garab, The fluid-mosaic membrane theory in the context of photosynthetic membranes: Is the thylakoid membrane more like a mixed crystal or like a fluid?. J. Plant Physiol. 252, 153246 (2020).

21. M. S. Webb, B. R. Green, Biochemical and biophysical properties of thylakoid acyl lipids. Biochim. Biophys. Acta Bioenerg. 1060, 133–158 (1991).

22. B. Fuks, F. Homble, Permeability and electrical properties of planar lipid membranes from thylakoid lipids. Biophys. J. 66, 1404–1414 (1994).

23. P. A. Armond, L. A. Staehelin, Lateral and vertical displacement of integral membrane proteins during lipid phase transition in Anacystis nidulans. Proc. Natl. Acad. Sci. U.S.A. 76, 1901–1905 (1979).

24. H. Wada, R. Hirasawa, T. Omata, N. Murata, The lipid phase of thylakoid and cytoplasmic membranes from the blue-green algae (cyanobacteria), Anacystis nidulans and Anabaena variabilis. Plant Cell Physiol. 25, 907–911 (1984).

25. W. Verwer, P.T Ververgaert, J. Leunissen-Bijvelt, A. J. Verkleij, Particle aggregation in photosynthetic membranes of the blue-green alga Anacystis nidulans. Biochim. Biophys. Acta Bioenerg. 504, 231–234 (1978).

26. Y. Tsukamoto, T. Ueki, T. Mitsui, T. A. Ono, N. Murata, Relationship between growth temperature of Anacystis nidulans and phase transition temperature of its thylakoid membranes. Biochim. Biophys. Acta Biomembr. 602, 673–675 (1980).

27. P. A. Siegenthaler, Molecular organization of acyl lipids in photosynthetic membranes of higher plants. In Lipids in photosynthesis: structure, function and genetics, 1st ed.; Springer, Dordrecht, 119–144 (1998).

28. S. Duchêne, P. A. Siegenthaler, Do glycerolipids display lateral heterogeneity in the thylakoid membrane?. Lipids 35, 739–744 (2000).

29. Y. Yamamoto, R. C. Ford, J. Barber, Relationship between thylakoid membrane fluidity and the functioning of pea chloroplasts: effect of cholesteryl hemisuccinate. Plant Physiol. 67, 1069–1072 (1981).

30. M. Velitchkova, D. Lazarova, A. Popova, Response of isolated thylakoid membranes with altered fluidity to short term heat stress. Physiol. Mol. Biol. Plants. 15, 43–52 (2009).

31. Z. Gombos, H. Wada, N. Murata, The recovery of photosynthesis from low-temperature photoinhibition is accelerated by the unsaturation of membrane lipids: a mechanism of chilling tolerance. Proc. Natl. Acad. Sci. U.S.A. 91, 8787–8791 (1994).

32. J. Li, L. N. Liu, Q. Meng, H. Fan, N. Sui, The roles of chloroplast membrane lipids in abiotic stress responses. Plant Signal Behav. 15, e1807152 (2020).

33. K. Gounaris, A. R. R. Brain, P. J. Quinn, W. P. Williams, Structural reorganisation of chloroplast thylakoid membranes in response to heat-stress. Biochim. Biophys. Acta Bioenerg. 766, 198–208 (1984).

34. I. Zaharieva, M. Velitchkova, V. Goltsev, Effect of cholesterol and benzyl alcohol on prompt and delayed chlorophyll fluorescence in thylakoid membranes. In Photosynthesis: Mechanisms and Effects, 1st ed.; Garab G., Ed.; Springer, Dordrecht 1827–1830 (1998).

35. M. Velitchkova, A. Popova, T. Markova, Effect of membrane fluidity on photoinhibition of isolated thylakoids membranes at room and low temperature. Z. Naturforsch C. 56, 369–374 (2001).

36. J. P. Douliez, A. Leonard, E. J. Dufourc, Restatement of order parameters in biomembranes: calculation of CC bond order parameters from CD quadrupolar splittings. Biophysi J. 68, 1727–1739 (1995).

37. K. Akabori, J. F. Nagle, Structure of the DMPC lipid bilayer ripple phase. Soft Matter. 11, 918–926 (2015).

38. P. Khakbaz, J. B. Klauda, Investigation of phase transitions of saturated phosphocholine lipid bilayers via molecular dynamics simulations. Biochim. Biophys. Acta Biomembr. 1860, 1489–1501 (2018).

39. F. de Meyer,B. Smit, Effect of cholesterol on the structure of a phospholipid bilayer. Proc. Natl. Acad. Sci. U.S.A. 106, 3654–3658 (2009).

40. J. A. Clarke, A. J. Heron, J. M. Seddon, R. V. Law, The diversity of the liquid ordered (Lo) phase of phosphatidylcholine/cholesterol membranes: a variable temperature multinuclear solid-state NMR and x-ray diffraction study. Biophysi J. 90, 2383–2393 (2006).

41. F. A. Heberle, G. W. Feigenson, Phase separation in lipid membranes. Cold Spring Harb Perspect. Biol. 3, a004630 (2011).

42. R. X. Gu, S. Baoukina, D. P. Tieleman, Phase separation in atomistic simulations of model membranes. J. Am. Chem. Soc. 142, 2844–2856 (2020).

43. M. Javanainen, H. Martinez-Seara, I. Vattulainen, Nanoscale membrane domain formation driven by cholesterol. Sci. Rep. 7, 1–10 (2017).

44. E. Giray et al., Multiscale simulations of biological membranes: the challenge to understand biological phenomena in a living substance. Chem. Rev. 119, 5607–5774 (2019).

45. M. Manna et al., Mechanism of allosteric regulation of β2-adrenergic receptor by cholesterol. Elife 5, e18432 (2016).

46. M. Manna, T. Nieminen, I. Vattulainen, Understanding the role of lipids in signaling through atomistic and multiscale simulations of cell membranes. Annu. Rev. Biophysics 48, 421–439 (2019).

47. M. Manna, et al., Long-chain GM1 gangliosides alter transmembrane domain registration through interdigitation. Biochim. Biophys. Acta Biomembr. 1859, 870–878 (2017).

48. C. S. Poojari, K. C. Scherer, J. S. Hub, Free energies of membrane stalk formation from a lipidomics perspective. Nat. Commun. 12, 1–10 (2021).

49. M. Manna, R. K. Murarka, Polyunsaturated fatty acid modulates membrane-bound monomeric α-synuclein by modulating membrane microenvironment through preferential interactions. ACS Chem. Neurosci. 12, 675–688 (2021).

50. H. J. Risselada, S. J. Marrink, The molecular face of lipid rafts in model membranes. Proc. Natl. Acad. Sci. U.S.A. 105, 17367–17372 (2008).

51. A. J. Sodt, M. L. Sandar, K. Gawrisch, R. W. Pastor, E. Lyman, The molecular structure of the liquid-ordered phase of lipid bilayers. J. Am. Chem. Soc. 136, 725–732 (2014).

52. A. J. Sodt, R. W. Pastor, E. Lyman, Hexagonal substructure and hydrogen bonding in liquid-ordered phases containing palmitoyl sphingomyelin. Biophysi J. 109, 948–955 (2015).

53. F. J. van Eerden, D. H. de Jong, A. H. de Vries, T. A. Wassenaar, S. J. Marrink, Characterization of thylakoid lipid membranes from cyanobacteria and higher plants by molecular dynamics simulations. Biochim. Biophys. Acta Biomembr. 1848, 1319–1330 (2015).

54. K. Ogata, T. Yuki, M. Hatakeyama, W. Uchida, S. Nakamura, All-atom molecular dynamics simulation of photosystem II embedded in thylakoid membrane. J. Am. Chem. Soc. 135, 15670–15673 (2013).

55. S. Vasil’ev, D. Bruce, A protein dynamics study of photosystem II: the effects of protein conformation on reaction center function. Biophysi J. 90, 3062–3073 (2006).

56. P. Kumari, CRK. Reddy, B. Jha, Comparative evaluation and selection of a method for lipid and fatty acid extraction from macroalgae. Anal. Biochem. 415, 134–144 (2011).

57. P. Kumari, study of lipids fatty acids and their derivatives in seaweeds. Shodhganga, Department of Biotechnology, Govt, of India, http://hdl.handle.net/10603/36894.

58. S. K. Mandal, V. R. Patel, G. Temkar, B. M. George, M. Raman, Bio-optic characterization of Discosphaera tubifer bloom occurs in an overcrowded fishing harbour at Veraval, India. Environ. Monit. Assess. 187, 1–16 (2015).

59. T. Róg, M. Pasenkiewicz-Gierula, I. Vattulainen, M. Karttunen, Ordering effects of cholesterol and its analogues. Biochim. Biophys. Acta Biomembr. 1788, 97–121 (2009).

60. S. Chakraborty et al., How cholesterol stiffens unsaturated lipid membranes. Proc. Natl. Acad. Sci. U.S.A. 117, 21896–21905 (2020).

61. F. Tardy, M. Havaux, Thylakoid membrane fluidity and thermostability during the operation of the xanthophyll cycle in higher-plant chloroplasts. Biochim. Biophys. Acta Biomembr. 1330, 179–193 (1997).

62. S. Munné-Bosch, The role of α-tocopherol in plant stress tolerance. J. Plant Physiol. 162, 743–748 (2005).

63. V. A. Mantri et al., An appraisal on commercial farming of Kappaphycus alvarezii in India: success in diversification of livelihood and prospects. J. Appl. Phycol. 29, 335–357 (2017).

64. A. Arasamuthu, J. K. Edward, Occurrence of ice-ice disease in seaweed Kappaphycus alvarezii at Gulf of Mannar and Palk Bay, Southeastern India. Indian J. Geomarine Sci. 47, 1208–1216 (2018).

65. J. B. Klauda et al., Update of the CHARMM all-atom additive force field for lipids: validation on six lipid types. J. Phys. Chem. 114, 7830–7843 (2010).

66. M. J. Abraham et al., GROMACS: High performance molecular simulations through multi-level parallelism from laptops to supercomputers. SoftwareX. 1, 19–25 (2015).

