## Supporting Information for "Phase transition in atomistic simulations of model membrane with thylakoid lipids of red algae"

**Method**

All simulations are performed with all-atom CHARMM36 force field parameters for lipids and salt, and CHARMM-modified TIP3P model for water (1-3). This set of parameters has been used in many recent computer simulations of lipid membranes (4-5). All simulations are conducted using GROMACS version 5.1.1 (6-7) and under an isothermal-isochoric (NPT) ensemble. A timestep of 2 fs was employed for integrating the equations of motion. Linear Constraint Solver (LINCS) algorithm (8) is employed to constrain the covalent bond lengths to hydrogen atoms. The v-rescale (stochastic velocity rescaling) thermostat (9) with a 1.0 ps coupling constant is employed to maintain the temperature of the system at seven different temperatures: 10 °C,15 °C, 20 °C, 25 °C, 30 °C, 35 °C, and 40 °C. The temperatures of the lipid membrane and solvent (water and ions) are controlled independently. The pressure of the system is controlled semi-isotropically at 1 bar using a Parrinello–Rahman barostat (10) with a 5.0 ps coupling constant. Periodic boundary conditions are applied in all three directions. The long-range electrostatic interactions are treated with the particle mesh Ewald (PME) method (11) with a real space cut-off of 10 Å. The van der Waals interactions are computed using the Lennard-Jones potential and using a cut-off distance of 10 Å. The neighbor lists are updated every 20 steps with a cut-off of 10 Å.

Both the solvated model bilayer membranes used in this study, Thylakoid-LBM and Chol-Thylakoid-LBM, are first energy minimized. Then a short simulation of 1 ns is performed with harmonic position restraints on lipid glycerol carbon atoms. Subsequently, all restraints are released and the system is subjected to production simulations at 7 different temperatures ranging from 10-40 °C. We have conducted 3 independent simulations of the membranes at a given temperature. For Thylakoid-LBM system each of the production simulations runs for 1600 ns, except simulations at 40 °C are 1000 ns long. For Chol-Thylakoid-LBM system production simulations are 1000 ns long. In presence of cholesterol the membrane properties, such as the area per lipid molecules, reaches plateau within 400 ns, while for in absence of cholesterol the area per lipid converges relatively slowly (Fig. S3). The first 400 ns of simulations for Chol-Thylakoid-LBM systems and first 800 ns for Thylakoid-LBM (except for simulations of Thylakoid-LBM at 40 °C, for which first 400 ns) are considered as equilibration period. Analyses are performed over the rest of the trajectory and averaged over independent trajectories at a given temperature (unless the results of individual trajectories are presented separately). The cumulative length of atomistic simulation used in this study is ~55 µs. VMD (12) was used for trajectory visualization.

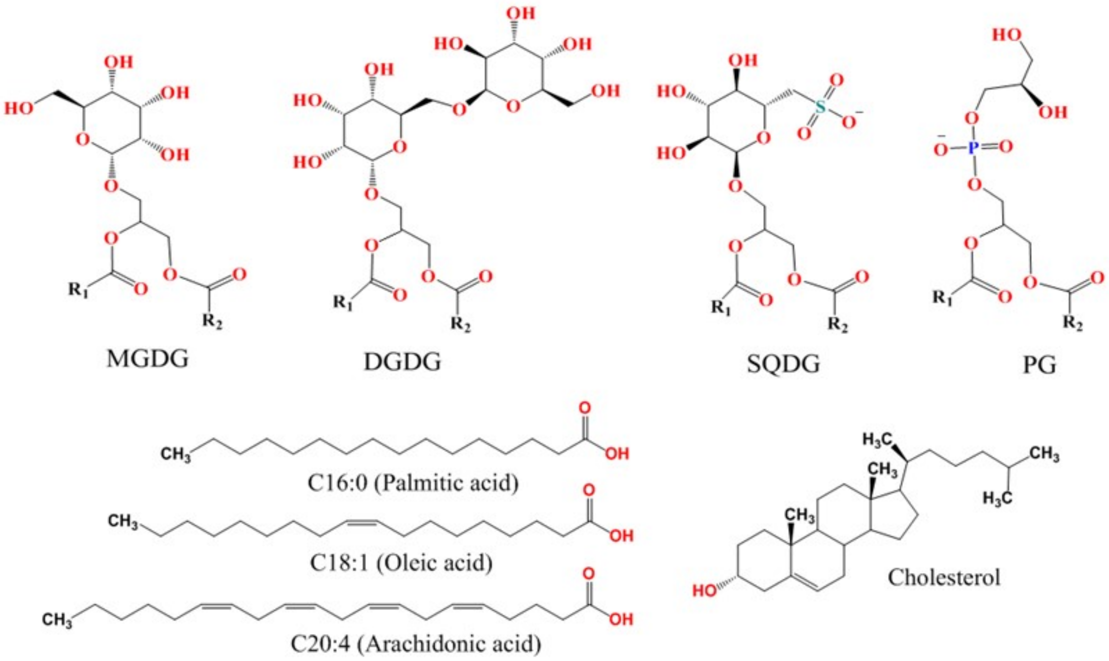


**Fig. S1.** The chemical structures of lipid head groups, fatty acid chains, and cholesterol.

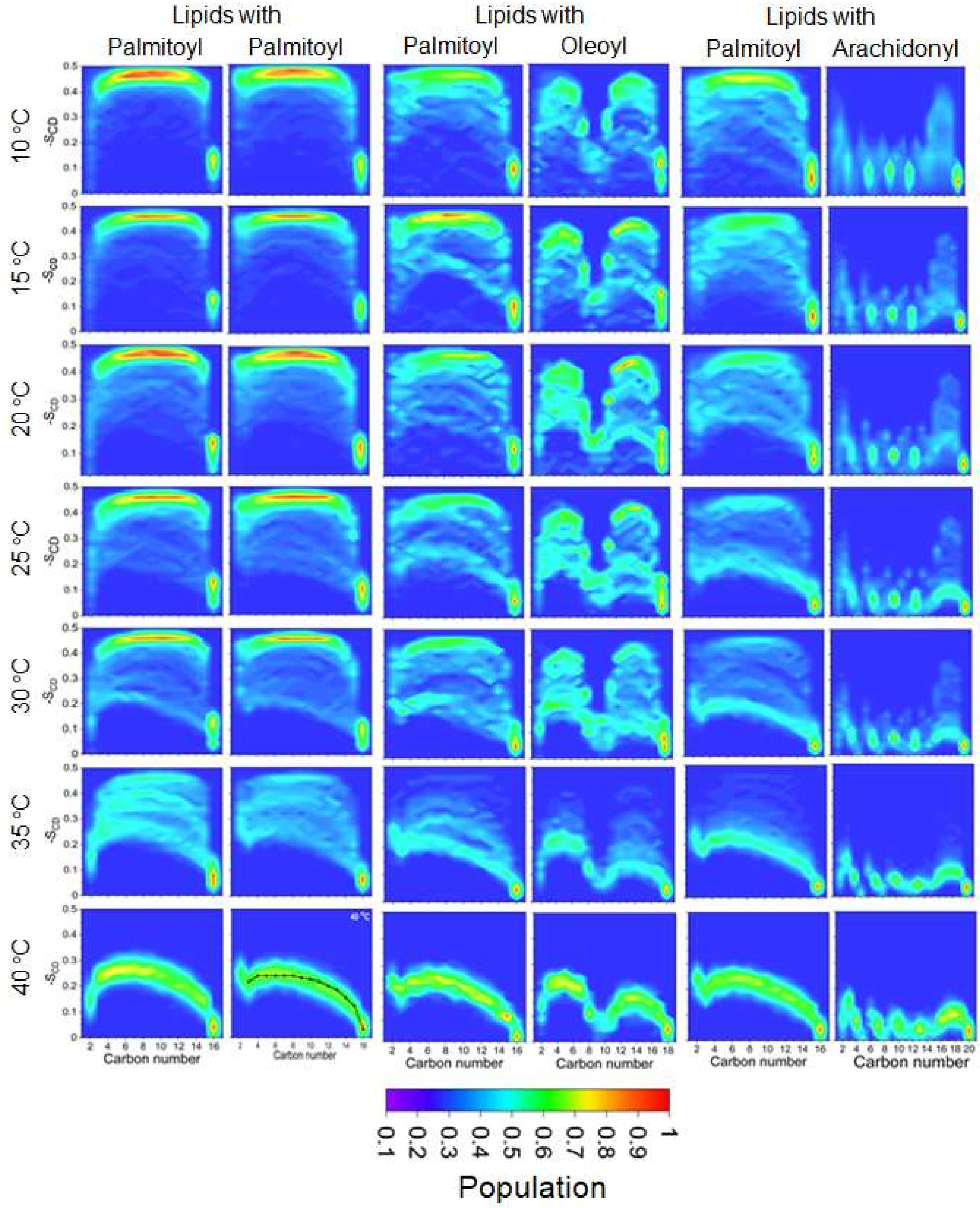


**Fig. S2.** Distribution of lipid acyl chain order parameter of the thylakoid membrane model as a function of temperature.

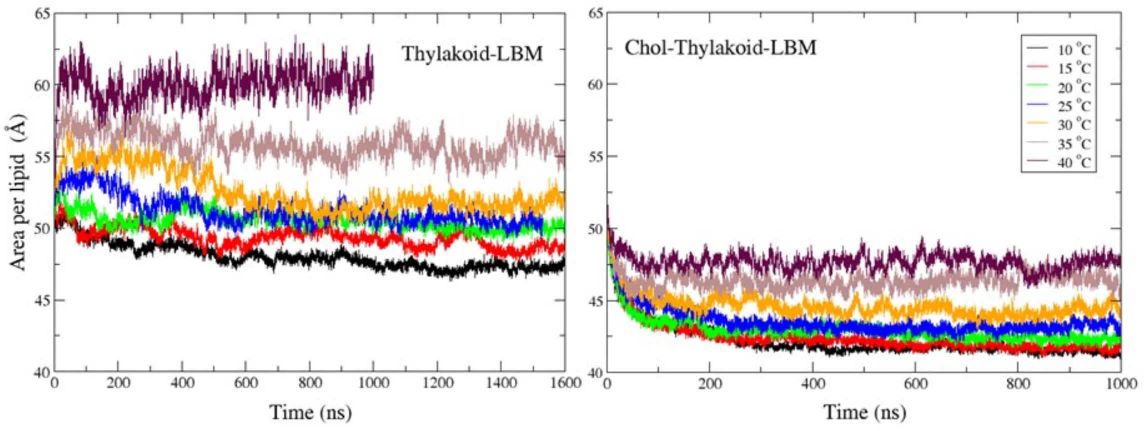


**Fig. S3.** Time profile of the area per lipid as a function of temperature.

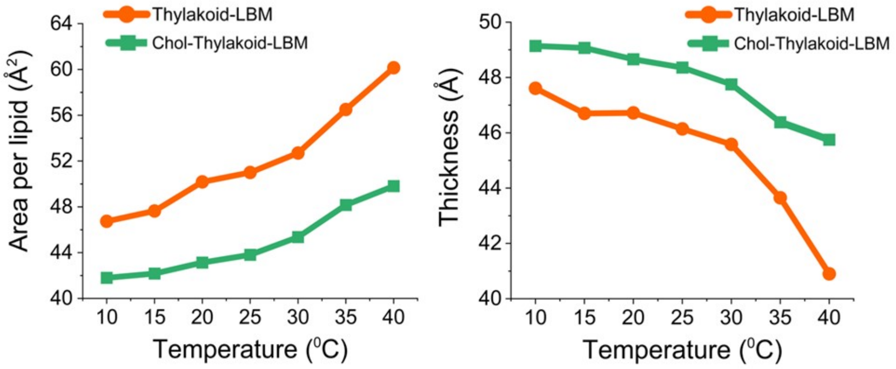


**Fig. S4.** Average area-per-lipid and average bilayer thickness as a function of temperature.

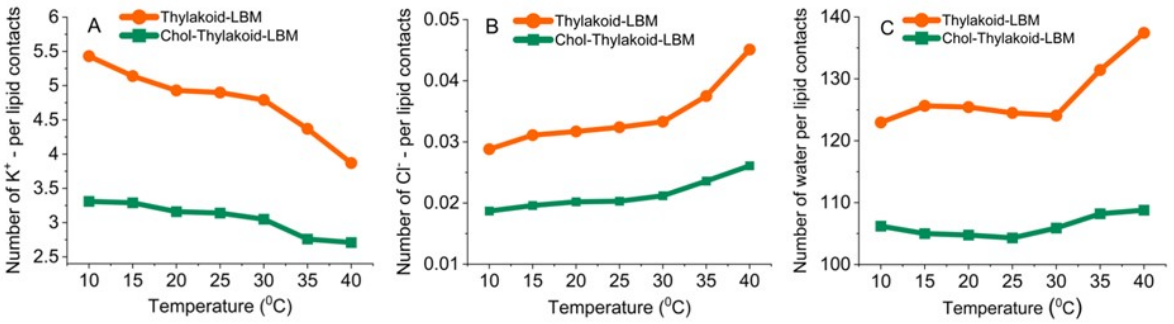


**Fig. S5.** Number of lipid-ion and lipid-water contacts as a function of temperature.

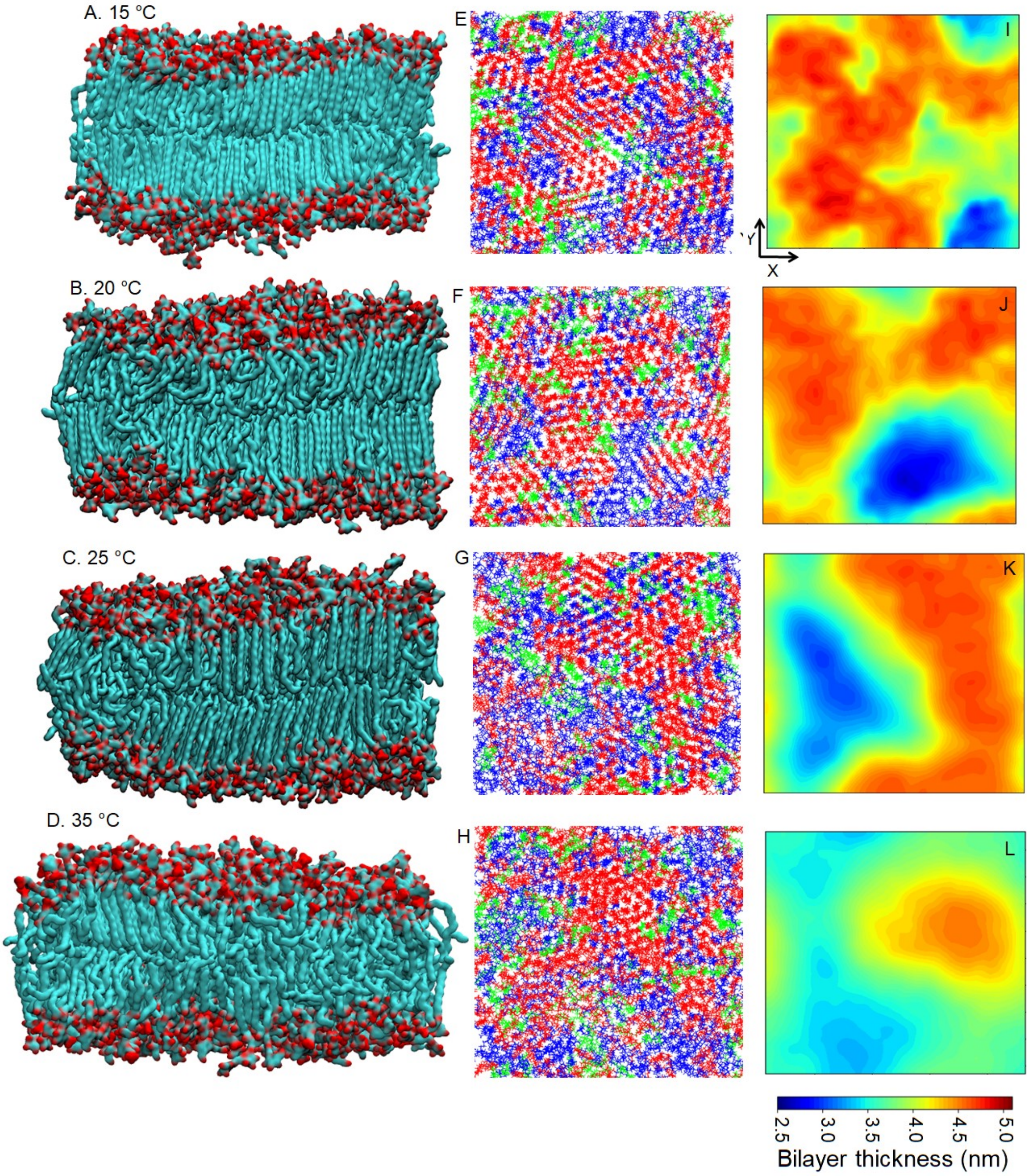


**Fig. S6.** Representative snapshots showing: (A-D) side-views and (E-H) top-views of the Thylakoid-LBM at different temperatures. The colour code is same as in Figure 4. (I-L) Bilayer thickness (nm) projected on membrane plane.

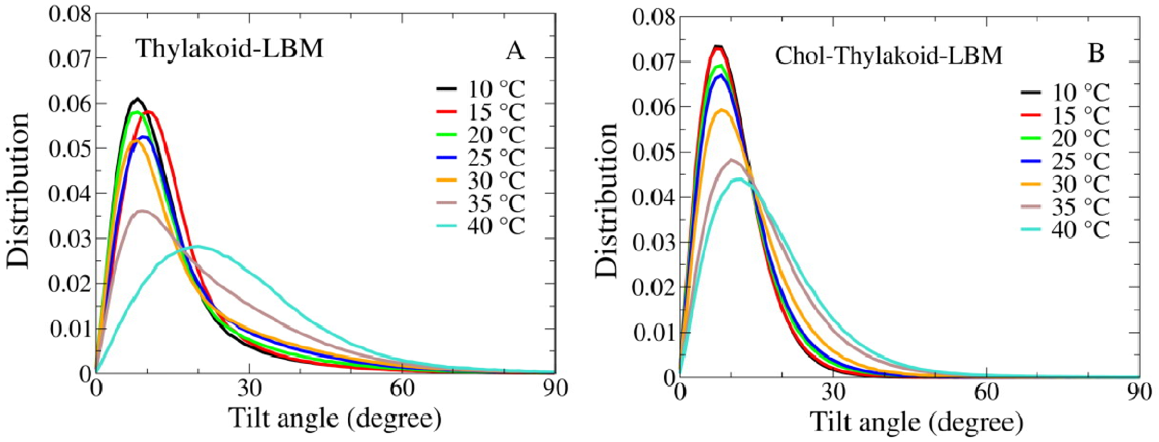


**Fig. S7.** Distributions of lipid acyl chain (sn-1 palmitoyl chain) tilt angle in Thylakoid-LBM and Chol-thylakoid-LBM at different temperatures.

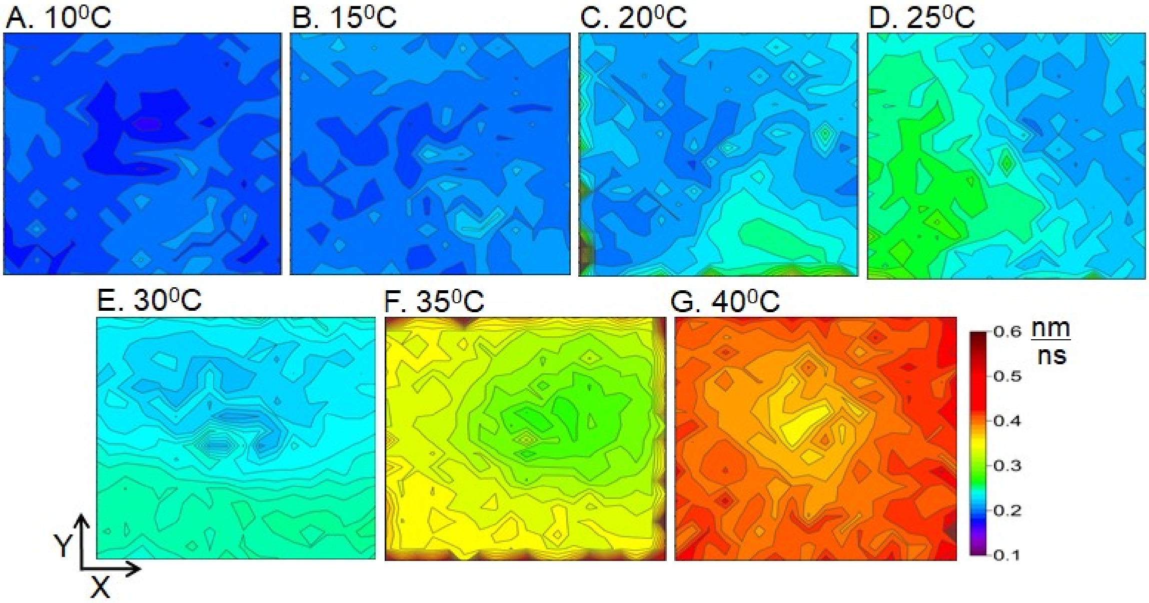


**Fig. S8.** In-plane lipid displacement (Jump map) in Thylakoid-LBM at different temperatures.

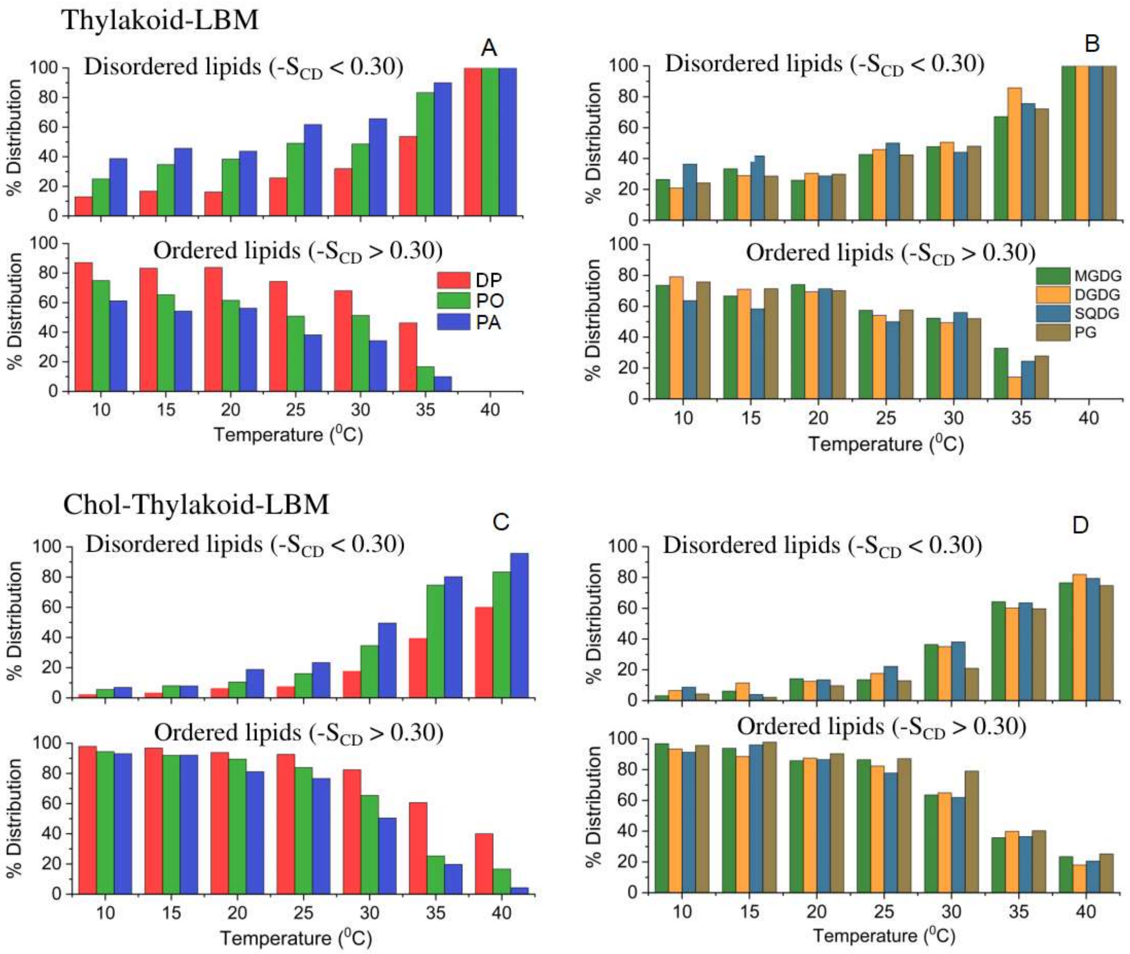


**Fig. S9.** Distribution of lipid acyl chain types (di-palmitoyl, DP; palmitoyl-oleoyl, PO; and palmitoyl-arachidonoyl, PA) and polar head group types (MGDG, DGDG, SQDG and PG) in ordered and disordered regions.

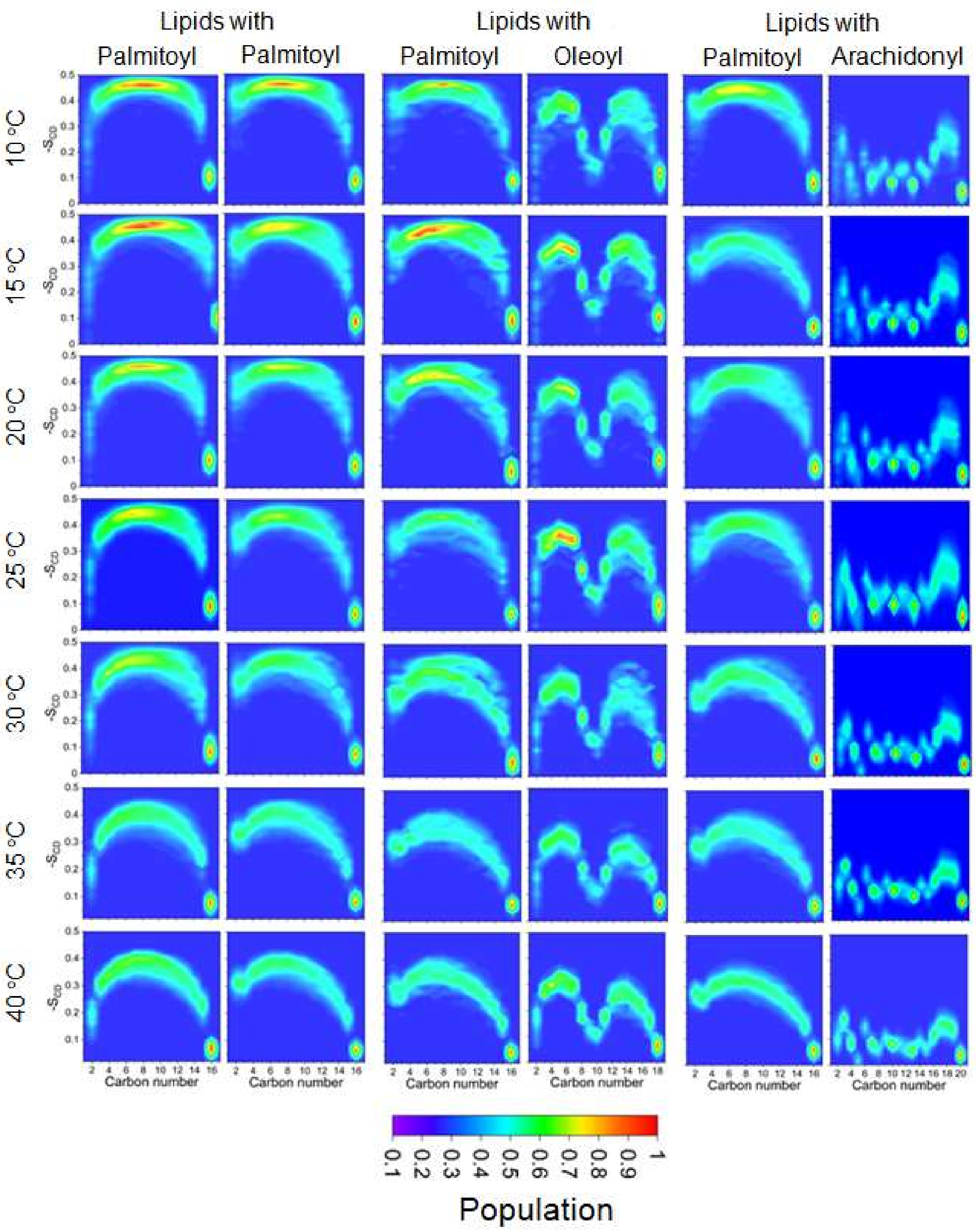


**Fig. S10.** Distribution of lipid acyl chain order parameter in Chol-Thylakoid-LBM as a function of temperature.

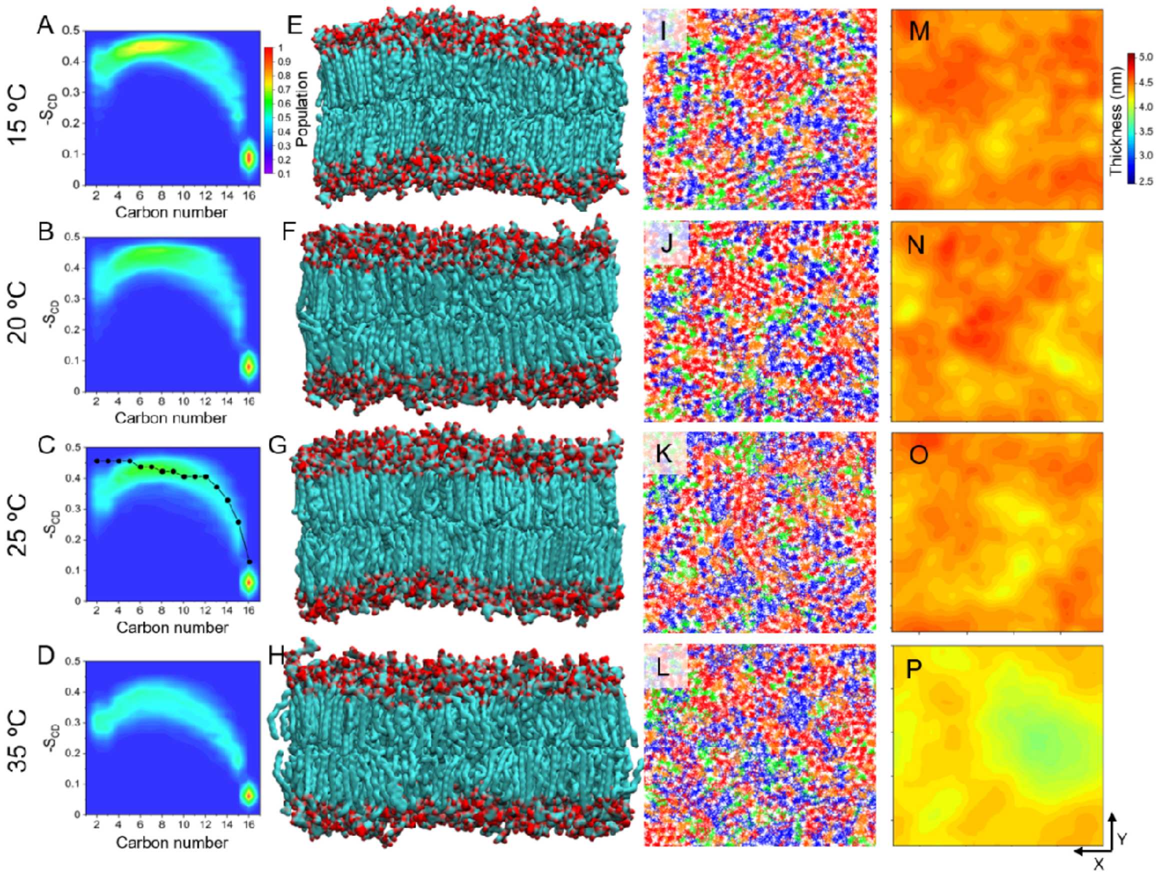


**Fig. S11.** Properties of Chol-Thylakoid-LBM. (A-D) -SCD distribution along carbon position of the sn-1 palmitoyl chain. The experimental -SCD profile from ref (47) is shown as black line with dots. Representative snapshots showing: (E-H) side-views and (I-L) top-views of the model membrane at different temperatures. The colour code is same as in Figure 4. (M-P) Bilayer thickness (nm) projected on membrane plane.

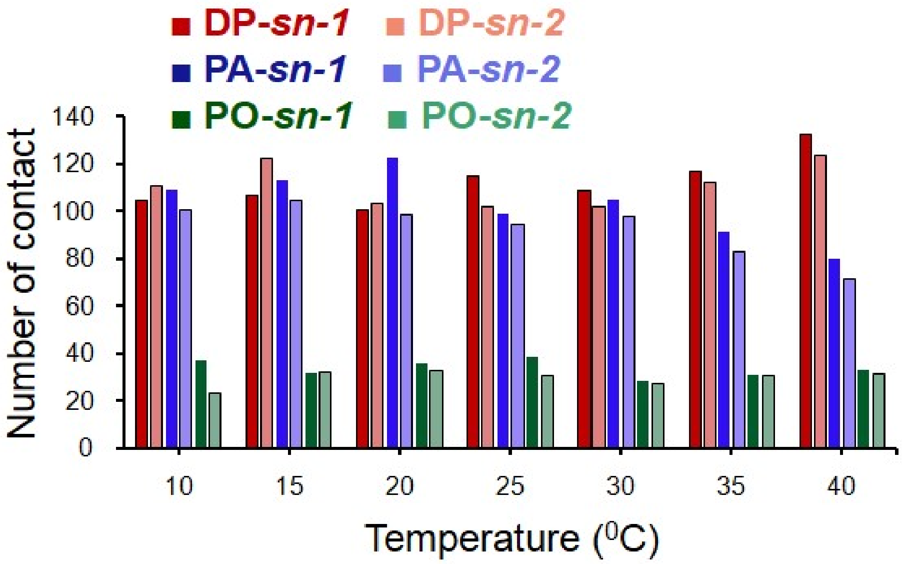


**Fig. S12.** Number of cholesterol-acyl chain contacts.

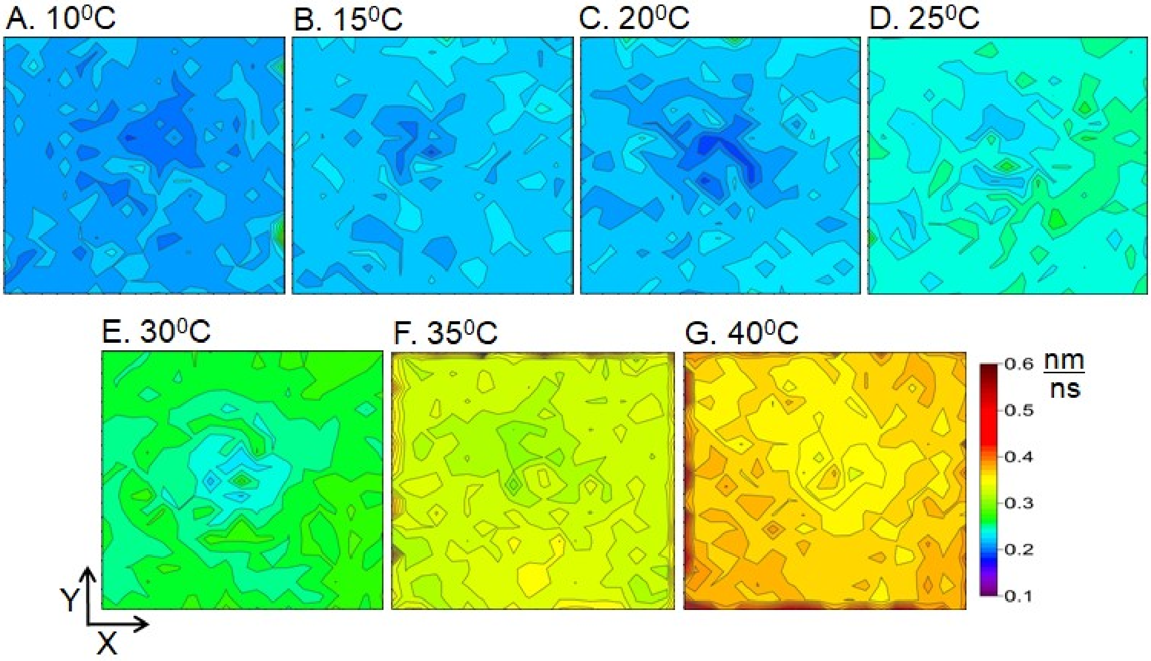


**Fig. S13.** In-plane lipid displacement (Jump map) in Chol-Thylakoid-LBM at different temperatures.

**SI References**


1. J. B. Klauda, V. Monje, T. Kim, W. Im, Improving the CHARMM force field for polyunsaturated fatty acid chains. *J. Phys. Chem. B* **116**, 9424-9431 (2012).
2. J. B. Klauda et al., Update of the CHARMM all-atom additive force field for lipids: validation on six lipid types. *J. Phys. Chem. B* **114**, 7830-7843 (2010).
3. W. L. Jorgensen, J. Chandrasekhar, J. D. Madura, R. W. Impey, M. L. Klein, Comparison of simple potential functions for simulating liquid water. *J. Chem. Phys.* **79**, 926-935 (1983).
4. M. Manna, R. K. Murarka, Polyunsaturated Fatty Acid Modulates Membrane-Bound Monomeric α-Synuclein by Modulating Membrane Microenvironment through Preferential Interactions. *ACS Chem. Neurosci.* **12**, 675-688 (2021).
5. Z. L. Li, P. Prakash, M. Buck, A "tug of war" maintains a dynamic protein–membrane complex: molecular dynamics simulations of C-Raf RBD-CRD bound to K-Ras4B at an anionic membrane. *ACS Cent. Sci.* **4**, 298-305 (2018).
6. D. Van Der Spoel et al., GROMACS: fast, flexible, and free. *J. Comput. Chem.* **26**, 1701-1718 (2005).
7. M. J. Abraham et al., GROMACS: High performance molecular simulations through multi-level parallelism from laptops to supercomputers. *SoftwareX* **1**, 19-25 (2015).
8. B. Hess, H. Bekker, H. J. Berendsen, J. G. Fraaije, LINCS: a linear constraint solver for molecular simulations. *J. Comput. Chem.* **18**, 1463-1472 (1997).
9. G. Bussi, D. Donadio, M. Parrinello, Canonical sampling through velocity rescaling. *J. Chem. Phys.* **126**,014101 (2007).
10. M. Parrinello, A. Rahman, Polymorphic transitions in single crystals: A new molecular dynamics method. *J. Appl. Phys.* **52**, 7182-7190 (1981).
11. T. Darden, D. York, L. Pedersen, Particle mesh Ewald: An N. log (N) method for Ewald sums in large systems. *J. Chem. Phys.* **98**, 10089-10092 (1993).
12. W. Humphrey, A. Dalke, K. Schulten, VMD: visual molecular dynamics. *J. Mol. Graph.* **14**, 33-38 (1996).
